# Visual speech enhances auditory onset timing and envelope tracking through distinct mechanisms

**DOI:** 10.1101/2024.11.23.624953

**Authors:** Cody Zhewei Cao, William C Stacey, Vibhangini S Wasade, Vernon L Towle, James X Tao, Shasha Wu, Naoum P Issa, David Brang

## Abstract

Seeing a speaker’s face facilitates speech recognition in challenging listening environments. Prior work has shown that visual speech contains timing information to aid auditory speech processing, yet how these signals are integrated within the auditory system during audiovisual speech perception remains poorly understood. Observation of preparatory mouth movements may initiate phase reset of intrinsic oscillations, potentially sensitizing the auditory system for receptive speech processing, while observation of mouth movements post speech onset may facilitate entrainment to the speech envelope. Yet, little work has been done to test whether visual speech enhances encoding of auditory speech onset, speech envelope tracking, or both, and through independent or overlapping mechanisms. To investigate this, we examined the ways in which visual speech timing information alters theta band power and phase using human intracranial electroencephalography (iEEG) recordings in a large group of patients with epilepsy (*n* = 21). Prior to speech onset, preparatory mouth movements elicited theta phase reset (increased inter-trial phase coherence; ITPC) throughout the superior temporal gyrus (STG), which is thought to enhance speech onset encoding. Following speech onset, visual speech modulated theta ITPC only at anterior STG electrodes while theta power was modulated at posterior STG electrodes. Pre- and post-speech onset were spatially and temporally dissociated, consistent with the hypothesis that audiovisual speech onset encoding and envelope tracking mechanisms are partially distinct. Crucially, congruent and incongruent visual speech, designed here to have identical visual timing information about speech onset time, but different visual mouth evolution, produced only a small difference in the phase of theta band oscillations in the anterior STG, highlighting a more restricted role of visual speech in ongoing auditory entrainment. These results support the hypothesis that visual speech improves the precision of auditory speech encoding through two separate mechanisms, with auditory speech onset encoded throughout the entire STG and ongoing speech envelope tracking within anterior STG.

## Introduction

In challenging listening environments, the presence of a speaker’s face facilitates speech recognition and comprehension (Sumby and Pollack 1959; Erber 1975; Ross et al., 2007), indicating that some information from the visual modality is relevant and beneficial to speech perception. Past research has examined what information is present in visual speech that aids speech perception processes, with studies indicating that visual timing information improves sensitivity to auditory speech onset (Abrams et al., 2008, Arnal et al., 2009, Karas et al., 2019) and entrainment to the speech envelope (for a review, see Obleser & Kayser,2019), that spectral information about the auditory signal can be recovered from mouth movements (Plass et al.,2020), and that lipreading can be used to bias phonemic processing (Karthik et al., 2024, Metzger et al., 2020). However, the independence of these processes, their neural mechanisms, and the functional roles of visual information on speech perception remain unclear.

Research has identified theta band oscillations as a candidate for how visual timing information influences auditory speech processes, with two potential mechanisms for brain-to-speech coupling. First, theta band oscillations are thought to encode the onset of auditory speech through phase-resetting mechanisms, leading to stronger oscillatory alignment in the theta band at speech onset (Giraud and Poeppel, 2012; Peelle and Davis, 2012; Ding and Simon,2014; Zoefel et al.,2018). Second, theta band power fluctuations correlate with the speech envelope, potentially reflecting enhanced time-locked processing of rhythmic speech information (e.g. Chalas et al., 2022, Ding et al. 2017; Keitel et al. 2018; Nourski et al., 2009). Thus, researchers have proposed that features of theta oscillations can encode both the onset of speech through phase-resetting mechanisms (onset encoding models) and the ongoing temporal dynamics of speech via sustained responses (envelope tracking models).

However, whether these two processes are supported by the same or distinct physiological processes remains a debate in the field. While some have argued that brain-to-speech coupling in theta band only encodes onset as part of an evoked response model (Oganian et al 2023), others have presented evidence that changes in theta oscillations throughout the speech envelope explains the brain-to-speech coupling equally as well as the evoked model in which only speech onset is encoded (Asama et al, 2024, for a review, see Duecker et al., 2024).

A parallel gap exists for understanding how visual speech timing information is used during speech perception. A large body of work has shown that audiovisual speech directly modulates activity in auditory areas, including the STG (e.g. Ghazanfar et al., 2005; Besle et al., 2008; Banks et al. 2011). Two separate mechanisms of audiovisual enhancements have been observed: first, in an evoked fashion, preparatory mouth movements prior to auditory speech onset can modulate auditory cortical responses through phase-resetting mechanisms in the theta band (Arnal et al., 2009; Luo et al., 2010; Mégevand et al., 2020). These visual-induced phase resets, as observed by higher ITPC prior to speech onset, are thought to facilitate precise timing predictions of upcoming auditory speech (Chandrasekaran et al., 2009; Schwartz & Savariaux, 2014; Karas et al., 2019). Second, in an entrainment fashion, as a word and sentence are heard, the observation of mouth movements increases the correlation between evoked neural responses in the auditory system and the speech envelope, which correlates with both subjective and objective measures of speech intelligibility (Ahissar et al. 2001, Crosse et al., 2015; O’Sullivan et al., 2017; O’Sullivan et al., 2020).

However, it remains unknown whether the same or different physiological processes in auditory cortex make predictions for speech onset based on preparatory mouth movements and track mouth evolutions during speech. Accordingly, the present study tests the hypothesis that different crossmodal mechanisms in the STG encode information about preparatory mouth movements before speech onset and mouth evolution during the speech envelope. We recorded intracranial EEG (iEEG) data from *n* = 21 epilepsy patients with electrodes implanted within the STG and presented single syllable words both in auditory-alone and audiovisual speech contexts. Importantly, the audiovisual speech condition included both congruent and incongruent audiovisual combinations (such that on some trials the visual signal mismatched the auditory signal) but with preserved onset timing information (i.e., the visual signal correctly predicted the onset time of the auditory word even in the incongruent condition). Using group-level linear mixed-effects modeling to compare auditory and audiovisual conditions, results showed that visual speech elicited a theta phase reset before auditory speech onset at the anterior, middle, and posterior STG. In parallel, we observed theta band power decreases in response to audiovisual speech after speech onset only in the posterior STG. Crucially, phase and power differences were dissociable at the group and individual subject levels, consistent with the hypothesis that visual speech encodes auditory onset and speech envelope through discrete mechanisms.

## Methods

### Participants, Implants and Recordings

*N* = 21 epilepsy patients (7 female, 30 ± 10.18-year-old) undergoing clinical evaluation using iEEG participated in this study. Data from *n* = 8 of these patients were included in a prior publication, but those analyses examined different features of the task data than those reported here (Karthik et al., 2024). iEEG data were recorded from clinically implanted depth electrodes (5-mm center-to-center spacing, 2-mm diameter) and/or subdural grid electrodes (10-mm center-to-center spacing, 3-mm diameter): 6 patients had subdural electrodes, and 18 patients had depth electrodes. Across all patients, data was recorded from a total of 1664 electrodes (83.2 ± 5.9 electrodes per participant, range = 30 - 170 per participant). The number, location, and type of electrodes used were based on the clinical needs of the participants. Due to differences between amplifiers used at different recording sites, iEEG data were obtained at 1000 Hz (*n* = 7), 1024 Hz (*n* = 2), or 4096 Hz (*n* = 12). All participants provided informed consent under Institutional Review Board (IRB) approved protocols at the University of Chicago (UC), University of Michigan (UM), and Henry Ford hospitals (HF). Single subject electrode distribution could be found in Supplemental Information Figure 1.

**Figure 1.**
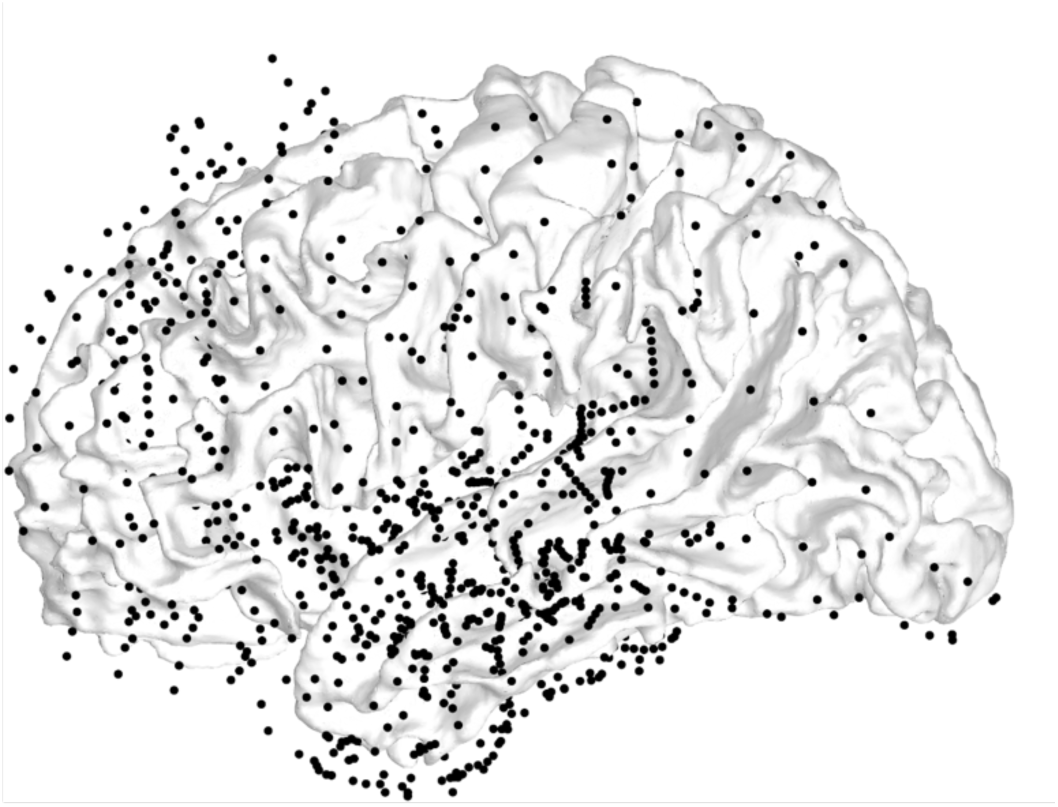
Electrode Distribution. All electrodes from *n*=21 patients who participated in the study. A total of *n*=1664 electrodes were planted in these patients, shown on the Freesurfer cvs_mni_152 white matter surface.

### iEEG Electrode Registration

Pre-operative T1-weighted magnetic resonance imaging (MRI) and a postoperative Computed Tomography (CT) scans were acquired for all participants in the study. Registration of the preoperative MRI to postoperative CT was performed using Statistical Parametric Mapping (Friston et al., 1994) and electrode localization was performed using custom software (Brang et al., 2016). This algorithm identifies and segments electrodes from the CT based on intensity values and projects subdural electrodes to the dura surface using the shape of the electrode disk to counteract post-operative compression. The Freesurfer image analysis suite (Dale, Fischl, and Sereno 1999; Fischl, Sereno, and Dale, 1999) was used for subsequent image processing procedures including cortical surface reconstruction, volume segmentation.

### iEEG Preprocessing

Data were resampled to 1024 Hz during initial stages of processing for all participants. To ensure that the observed signals were analyzed from maximally local neuronal populations, data were referenced offline in a bipolar fashion (signals subtracted from each immediately adjacent electrode in a pairwise manner). Noisy electrodes were identified and rejected by estimating raw signal variance. Channels in which variability exceeded 5 SD were rejected. Data were then split into 2-second epochs (−1 to 1 seconds pre-stimulus time with t = 0 at the onset of auditory speech). Noisy trials were likewise identified and rejected when variability in the raw signal exceeded 3 SD. (across all trials within the electrode). Slow drift artifacts and power-line interference were attenuated by high-pass filtering the data at 0.1 Hz and notch-filtering at 60 Hz (and its harmonics at 120, 180, and 240 Hz). Individual trials were then separately filtered into theta (3 - 7 Hz, wavelet cycles varied linearly from 3-5) using Morlet wavelet convolution and then power transformed.

### Electrode Selection

Electrodes meeting both anatomical and functional criteria were included in analyses. Electrodes met the *anatomical selection criteria* if they were located within 10 mm of the FreeSurfer anatomical label “superiortemporal”, derived from the MNI152 template (Collins et al., 1994; Fonov et al., 2011). Electrodes met the *functional selection criteria* if they produced a significant response in the theta band across all conditions in the [0, 500ms] time window, meaning that we observed a significant peak in the power spectral density plot (PSD) the theta band range (4-7 Hz). Among the electrodes that met the criteria for the anatomical localizer, a total of *n*=527 electrodes met the *functional* criteria, and were included in the study (see Table 1).

**Table 1.** Number of electrodes and participants who contributed data to each group-level time series analysis

| Anterior STG |  | Middle STG |  | Posterior STG |  |
| --- | --- | --- | --- | --- | --- |
| N. Elecs | N Subs | N. Elecs | N Subs | N. Elecs | N Subs |
| 176 | 18 | 217 | 19 | 134 | 18 |

### Auditory-Visual Speech Paradigm

Audiovisual speech stimuli were modified from a prior study (Ross et al., 2007) in which a female speaker produced common monosyllabic words. The modified stimulus set contained 40 common monosyllabic words spoken by a female speaker. Each word began with one of four initial consonants: /b/, /f/, /g/, and /d/ (10 of each); phonemes in the second position of these words were additionally balanced across the four groups. Stimuli were recorded at 29.97 frames per second, trimmed to 1100 ms (or 33 frames) in length, and adjusted so the first consonantal burst of sound occurred around *t=*500 ms during each video.

Patients were seated in a bed at the testing hospital (UM, HF & UC) where they performed a word identification task while auditory, visual, and audiovisual speech stimuli were presented via a laptop using Psychtoolbox (Brainard, 1997; Pelli, 1997). Subjects were presented with trials in one of four main conditions: auditory-alone, visual alone, congruent audiovisual or incongruent audiovisual. Each main condition had 40 trials per block for a total of 160 trials per block, and all subjects performed 2 blocks (320 trials total, 80 trials per main condition group). Half of the trials in each condition were presented as they were recorded, and in half, pink noise was added to the auditory channel to reduce the signal-to-noise (SNR) ratio of the signals to −6 db SPL. In the current study, trials were analyzed without consideration of the clear and noisy distinction.

Two task variants were used. In Task Variant 1, on each alternate trial, subjects (*n* = 3) were asked to speak aloud the last two words presented. In Task Variant 2 (see Fig. 2), subjects (*n* = 18) were asked to indicate the letter(s) associated with the initial sound of each word in a 4-alternative forced choice task. A pilot study was performed to validate stimuli use and study design where undergraduate students at the University of Michigan performed free response reports of the word they heard (Ahn et al., 2024). Based on piloting results, two sets of choices were available for subjects depending on the word identity. For trial stimulus beginning with a consonant of “D” or ‘F’, like ‘FISH’, response options could be ‘D’, ‘F’, ‘T’, and ‘TH’, corresponding to ‘DISH’, ‘FISH’, ‘TISH’, and ‘THISH’. For trial stimulus beginning with a consonant of “B” or ‘G’, like ‘BUY’, response options would be ‘B’, ‘G’, ‘D’, and ‘TH’, corresponding to ‘BUY’, ‘GUY’, ‘DIE’, and ‘THIGH’. Participants responded using a joystick device. The full stimuli list can be found in Supplemental Information(SI) Table 1.

**Figure 2.**
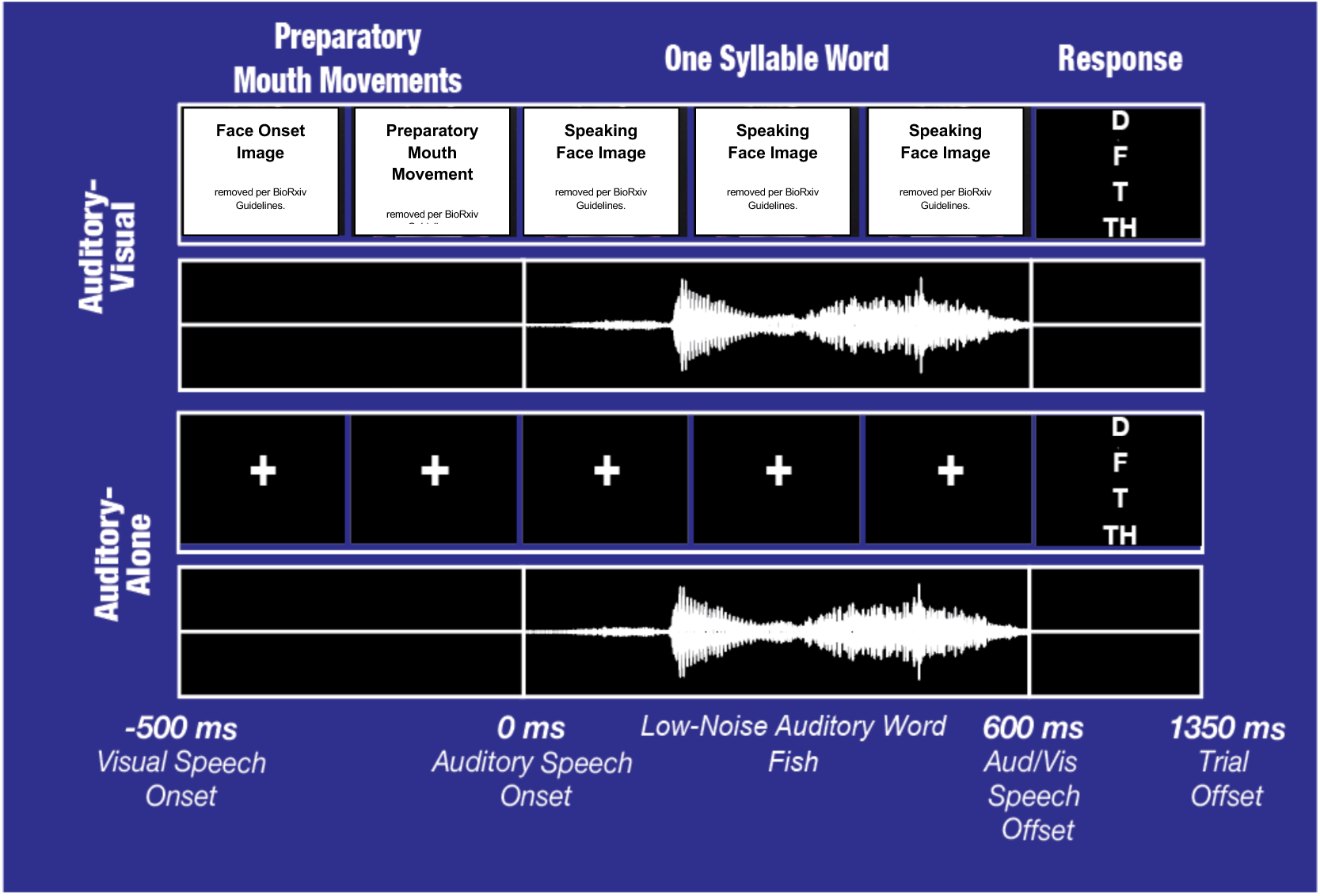
iEEG Experiment Paradigm. The current study adopted a 4 × 2 design in which the task conditions orthogonally manipulated (1) the visual speech information on a trial (no visual [auditory-alone], no auditory [visual-alone], congruent AV and incongruent AV), and (2) the intelligibility of the auditory speech stimuli (low-noise or high-noise). As shown in Figure 1, subjects were presented with words and were asked to select the initial consonant heard from four options in the current trial. For example, on a trial with the auditory word “FISH”, the options presented to the participant are the letters “D”, “F”, “T”, and “TH”, which correspond to the words “DISH”, “FISH”, “TISH”, and “THISH”.

## Analysis

### Group-Level Time-Series Analyses

We used linear mixed-effects models to estimate general group-level differences between auditory-only and audiovisual conditions as done previously by our group (e.g., Karthik et al. 2021 and Karthik et al. 2024) and in a similar framing to other groups (Desai et. al., 2024). We created three regions of interest (ROIs) within the STG to provide coarse anatomical delineation while ensuring an adequate amount of data was present in the ROIs to fit the LME models. ROIs were divided into three equal partitions from the “superiortemporal” label in Freesurfer, comprising anterior, middle, and posterior regions, similar to the division of the STG used previously (Smith et al., 2013). Electrodes within 10 mm of these labels were linked to the closest of the three. No electrode was assigned to more than one ROI.

Linear mixed-effects modeling of the time-series data was performed using the fitlme function in MATLAB R2021a (MathWorks Inc., Natick, MA). First, electrodes in the same ROI from the same participant were averaged at each timepoint. The justification for the averaging step is two-fold: one, neighboring electrodes share variance in the model, and two, to reduce complexity of the model. Two, individual trial theta power time-series were down-sampled to 100Hz, and trimmed to the [-1, 0.5] second window relative to auditory speech onset, leaving a total of 151 data points per trial per ROI per subject. Lastly, main-effect models were constructed at each of the 151 timepoints, in which differences between auditory-only and audiovisual trials, or congruent-audiovisual and incongruent-audiovisual conditions, were separately evaluated at each of the three STG ROIs (anterior, middle, posterior). In Wilkinson notation (Wilkinson & Rogers, 1973), the linear mixed effect model is represented as:

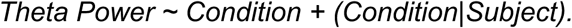

Critically, we modeled both random intercepts and random slopes for trial conditions to maintain ‘maximal’ models for confirmatory hypothesis testing (Barr et al., 2013). Statistics for the main-effect models were adjusted for comparisons at multiple time-points from −500 to 500 ms using FDR correction (*q* = .05) (Groppe, Urbach, and Kutas, 2011).

### Analysis of Inter-Trial Phase-Coherence (ITPC)

For group-level analyses of transient inter-trial phase-coherence (ITPC), each epoch was convolved with 20 wavelets with center frequencies ranging from 1 to 20 Hz in 1-Hz intervals, with the number of cycles varying across this frequency range (3 cycles for the frequencies 1–3 Hz and 4–20 cycles varied linearly for the frequencies 4–20 Hz). For group-level comparisons against baseline, ITPC was calculated separately for each frequency band and each electrode during the pre-stimulus baseline period (−1,000 to −500 ms) and the 500 ms stimulus onset or offsets period (0–500 ms); we used a 500-ms window in the baseline period to ensure stable “null” ITPC values. Frequency-specific ITPC values for each period were subtracted and Fisher’s Z-transformed before averaging across frequencies to produce electrode-specific difference scores. These difference scores were then averaged across each participant’s auditory or visual electrodes, and these participant-level difference scores were compared to 0 using a one-tailed, one-sample *t*-test.

For group-level time-frequency analyses, ITPC values for each timepoint (−1 to 1 s) in the theta frequency band were first averaged across electrodes within each participant and then averaged across participants. Significant timepoints were identified in the −0.5 to 0.5 s range relative to the pre-stimulus baseline period (−1 to −0.6 s) using one-tailed one-sample *t* tests at each timepoint and frequency. Multiple comparison corrections were applied using cluster statistics, in which the time-frequency map of *t* value statistics is permuted 10,000 times to identify clusters of significant contiguous time-frequency points (at *p* < 0.05) that are greater than those in 95% of the permuted data.

For single-electrode analyses of transient ITPC, phase-shuffling was used to generate a null distribution of ITPC values for the 200-ms onset period in each electrode. For 10,000 iterations, a random value ranging from −π to +π (uniformly sampled) was added to each trial before calculation of ITPC. The distribution of ITPC values produced by this procedure is designed to reflect the expected distribution of values observed under purely spurious phase coherence. *p* values were determined by identifying the proportion of values from this distribution that equaled or exceeded observed ITPC values, and then were corrected for multiple comparisons using FDR correction (*q* = 0.05).

### LME Interaction at the Group Level

To test whether the phase and power modulations are independent, we modeled the interaction between congruent-audiovisual and auditory-alone conditions, for theta ITPC values in the −0.5 to 0 sec window (the preparatory mouth movement period), and theta power in the 0 to 0.5 sec window (the duration of auditory speech). The same time series data from congruent-audiovisual and auditory-alone conditions were used in a linear mixed-effects model to test for interaction between the two mechanisms separately at each of the three ROIs:

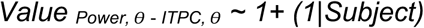

### Correlations at the Single Subject Level

To test for the presence of a correlation between phase and power metrics, a linear mixed-effects model, which accounts for inter-subject variability by adding subject as a random effect, was constructed to determine if there is a consistent pattern across participants. Time-series data from the theta phase and power analyses were modeled with the difference between the conditions as the fixed effect and participants as the random effect:

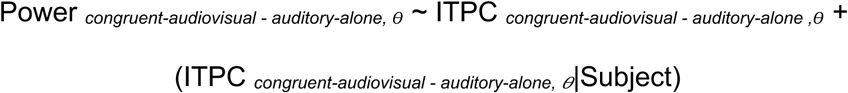

## Results

### Spectral Responses Across Conditions

We first examined theta band power across all electrodes to replicate findings of theta power increase following speech onset. Using a linear mixed effect model with subjects as the random effect with all electrodes throughout the STG (*n*=527) we quantified theta band power (4 - 8 Hz) after sound onset (0 ∼0.5 sec) relative to the pre-stimulus baseline. Consistent with prior findings (e.g., Karthik et. al. 2021), we found a significant increase in theta band power at the group level in each of the three conditions (Figure 3): auditory-alone (*F*(1,26876)=16.467, *p<*0.001); congruent-audiovisual (*F*(1,26876)=7.07, *p=*0.007), incongruent-audiovisual (*F*(1,26876)=6.182, *p*= 0.017).

**Figure 3.**
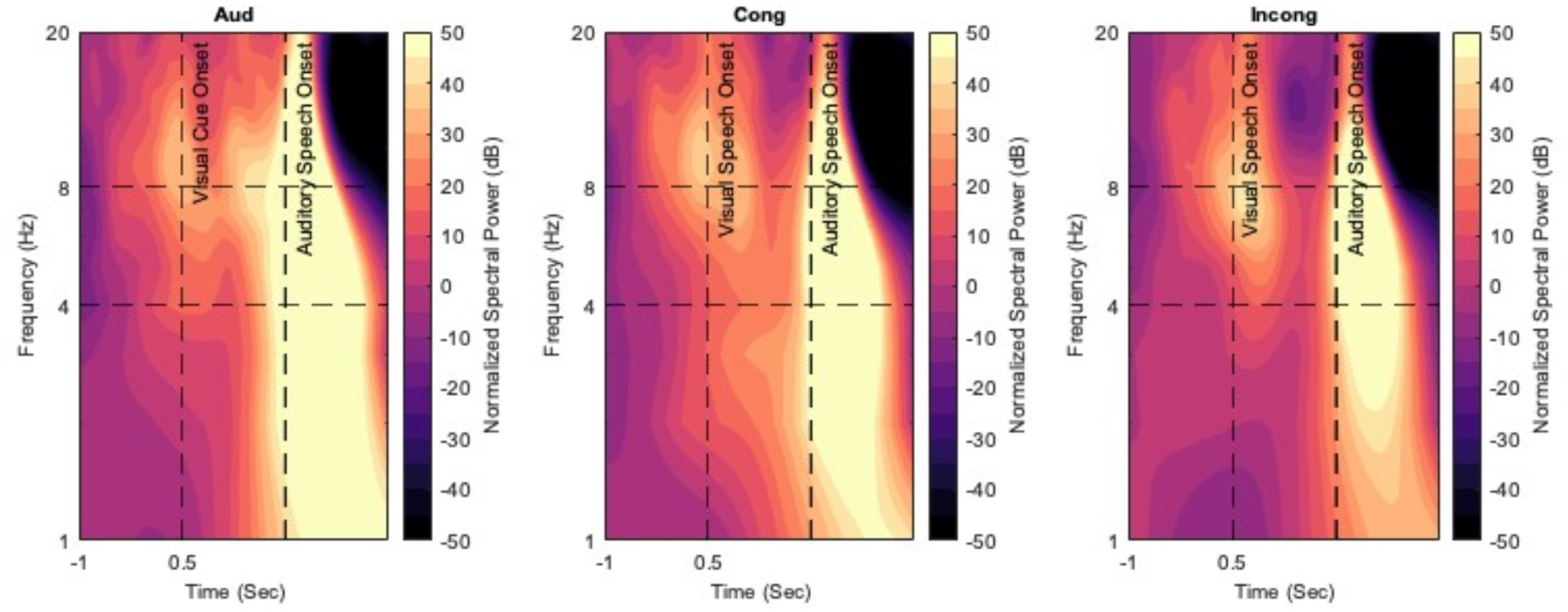
Group-level plots showing event-related spectral power from 1 to 20 Hz (shown in log scale), from −1 to 0.5 secs, for all three conditions. Data reflect iEEG activity from all anatomically localized auditory electrodes (n=501), first averaged across electrodes within each participant, then averaged across participants. Vertical dotted lines denote visual mouth movement onset (−0.5 sec) and auditory speech onset (0 sec). Theta band (4-8 Hz) is highlighted by two horizontal lines. Color scale reflects normalized power.

### Comparing Theta ITPC and Theta Power in Auditory-Alone and Congruent-Audiovisual Trials

First, we sought to replicate the finding that preparatory mouth movements elicit a theta phase reset (Luo et al., 2010; Mégevand et al., 2020; Biau et al., 2021). To do this, we compared the time courses of inter-trial phase-coherence (ITPC) values for both auditory-alone and congruent-audiovisual conditions in auditory electrodes across the STG. This analysis was performed by pooling electrodes within the anterior, middle and posterior STG within each subject (Smith, 2013) and modeled both the slope and intercept using a linear mixed-effects model.

Average ITPC responses from each condition and in each ROI are shown in Figure 4A. Prior to visual mouth movement onset, between −1 and −0.5 seconds, ITPC values for both congruent-audiovisual and auditory-alone remained non-significant from baseline throughout the STG. After mouth movement onset at −0.5 seconds, ITPC values in the congruent-audiovisual condition increased, peaking shortly after auditory speech onset, while ITPC values in the auditory-alone condition failed to increase until auditory speech onset. The conditional ITPC differences between congruent-audiovisual and auditory-alone were strongest during the preparatory mouth movement window in all three segments of the STG. Significantly greater ITPC (*p* < .05, FDR-corrected) was observed in the congruent-audiovisual condition compared to the auditory-alone condition across broad time ranges: anterior STG (−0.31 to 0.06 secs, and 0.42 to 0.50 sec, minimum *p* = 0.0269), middle STG (−0.45 to −0.09 sec, minimum *p* < 0.001) and posterior STG(−0.37 to −0.21 sec, minimum *p* = 0.002).

Second, we examined theta power changes across the two conditions. Prior research has demonstrated that following speech onset, congruent audiovisual trials elicit decreased theta power relative to auditory alone trials (Karthik et al., 2021). Accordingly, we compared the time courses of theta band power for both auditory-alone and congruent-audiovisual conditions in auditory electrodes across STG. As in the ITPC analyses, we pooled electrodes within each ROI per subject and modeled both the slope and intercept using a linear mixed-effects model.

Results showed a similar pattern to those in Karthik et al. (2021), a study using syllable-level stimuli. Theta power within all ROIs of the STG increased steadily before auditory onset and peaked immediately after sound onset (Figure 4B). Consistent with previous results using audiovisual phonemes (Karthik et al., 2021), less theta power was observed in the congruent-audiovisual condition compared to the auditory-alone condition generally after auditory onset, with effects only present in the posterior superior temporal gyrus (−0.08 to 0 sec and 0.24 to 0.50 sec, minimum *p*=0.0026). No significant differences were observed after correcting for multiple comparisons at the anterior STG (minimum *p =* 0.557) and middle STG (minimum *p = 0*.383).

**Figure 4.**
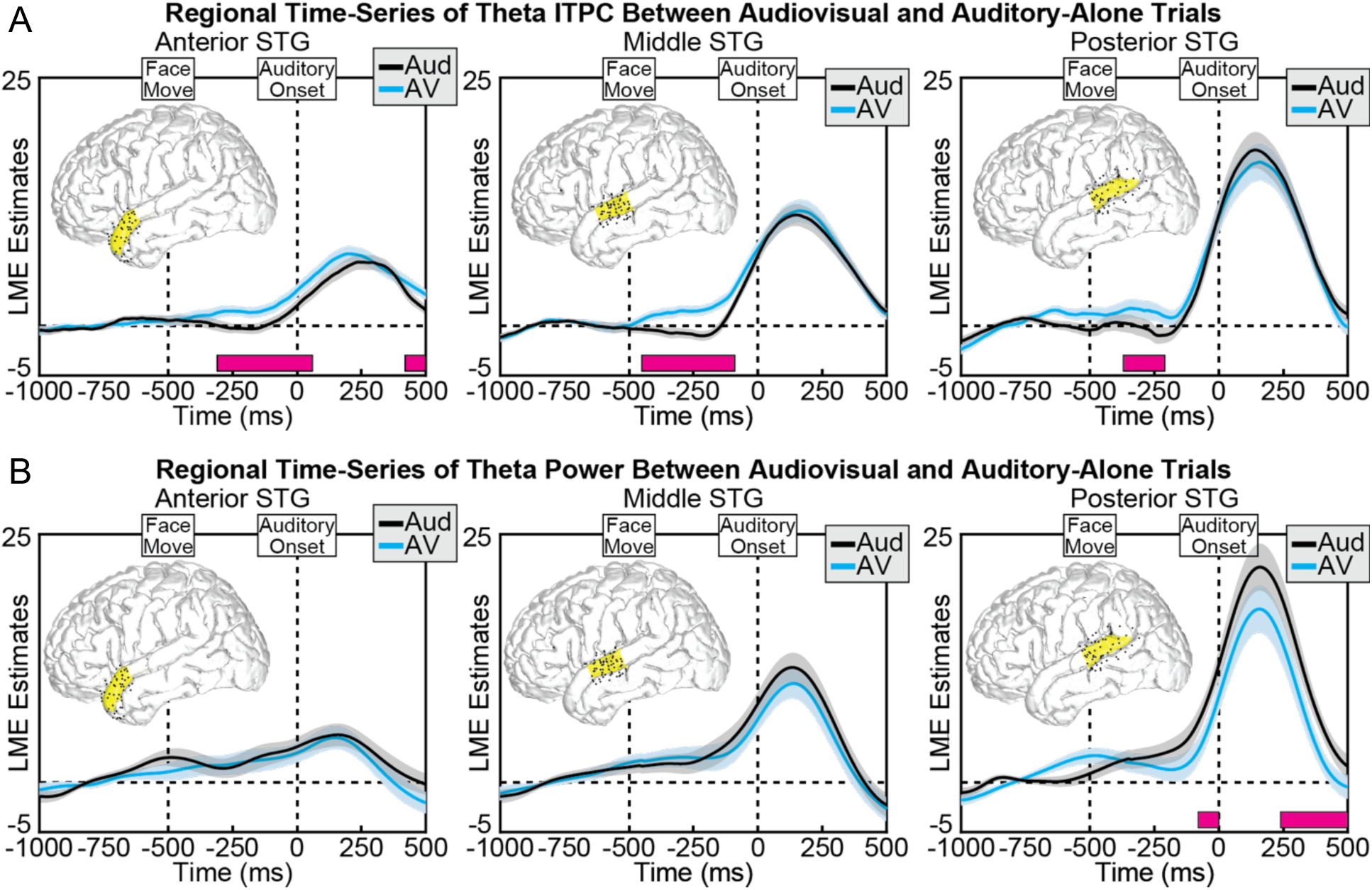
(a) Group linear mixed-effect model (LME) estimates for each time-point of theta ITPC in auditory-alone (blue) and audiovisual (red) trials, calculated separately at anterior (left), middle (middle), and posterior (right) regions of the STG. Shaded areas reflect 95% confidence intervals. Black boxes reflect significant differences after correcting for multiple comparisons. Significant differences in theta power occurred after visual speech onset in anterior, middle and posterior STG. (b) LME estimates for each time-point of theta power in auditory-alone (blue) and audiovisual (red) trials, calculated separately at anterior (left), middle (middle), and posterior (right) regions of the STG. Shaded areas reflect 95% confidence intervals. Black boxes reflect significant differences after correcting for multiple comparisons. Significant differences in theta power occurred mainly in the posterior STG.

### Interactions between Measures and Conditions

The present results showed two effects of visual speech on theta band activity. First, congruent-audiovisual trials showed increased theta ITPC relative to auditory-alone trials before speech onset throughout the STG (though largest in the middle STG). Second, congruent-audiovisual trials showed decreased theta power relative to auditory-alone trials generally after speech onset and only within the posterior STG. However, these results in isolation do not directly test whether the same or different mechanisms contribute to conditional differences before and after speech onsets.

Instead, if these pre-auditory-onset and post-auditory-onset effects are independent as we hypothesize, then we would expect to see a significant interaction between condition (congruent-audiovisual vs auditory-alone) and measure (ITPC vs Power). To test this, we calculated conditional differences for each measure at the electrode level using linear mixed effect models with subject as the random factor. Results showed significant interactions between measure (ITPC and power) and condition (congruent-audiovisual and auditory-alone) in each of the three STG regions: anterior STG [*F*(175)= 7.76, *p*=0.006], middle STG [*F*(216)=14.788, *p*<0.001], and posterior STG (*F*(133)= 24.92, *p*<0.001). The significant interaction observed between condition and measure at the electrode level suggests that different mechanisms contribute to effects before and after speech onsets.

### Correlations at the Individual Subject Level

The above test examined the relationship between the ITPC and power effects at three regions of the STG using a linear mixed effect model, showing a significant interaction between the two. However, this significant interaction does not preclude the possibility that both ITPC and power effects are still present at the same electrodes within a participant, which would run counter to our hypothesis that the effects arise from different neuronal populations. To test for a linear relationship between these effects, we calculated correlations between the ITPC and power differences at the single subject level using all electrodes within the STG) at the single subject level. Specifically, if it is true that these effects are generated by the same underlying mechanism, then we would expect a consistent pattern of negative correlations, observed at the single subject level. For example, an electrode showing a large positive effect for the ITPC analysis (i.e., ITPC_*θ*, congruent-audiovisual_ > ITPC _*θ*, auditory-alone_ before speech onset) should also show a large negative effect for the power analysis (i.e., Power _*θ*, congruent-audiovisual_< Power _*θ*, auditory-alone_ after speech onset). Consistent with our prediction that the phase and power effects are generally unrelated at the electrode level, only 1 out of 21 participants showed a significant correlation between the two measures (FDR-correction applied). The scatter plots of individual-subject correlations are shown in Figure 5.

**Figure 5.**
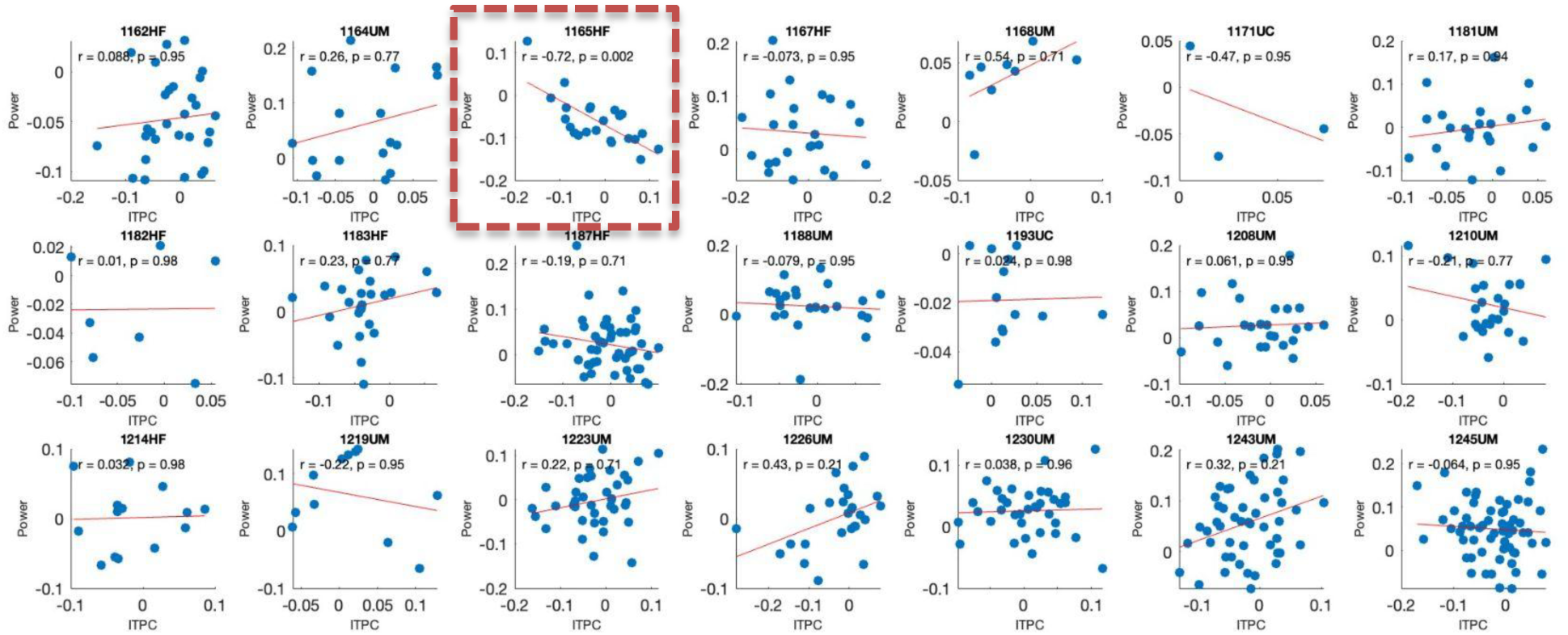
Single subject correlations of theta band ITPC differences between congruent audiovisual and auditory alone conditions during preparatory mouth movement period and theta band power differences between the same two conditions following speech onset. Each blue dot represents a single electrode in the STG included in the analysis. A linear regression (red line) between the two difference values was fitted at the individual subject level to quantify the relationship between the two processes. 20 out of 21 subjects returned no significant correlation between the two metrics (p>.05 after FDR correction). One subject showed a significant relationship (*r* = −0.72, *p* = 0.002), which is marked in red.

### Correlations at the Group Level

The individual participant correlation analyses showed that only one out of the 21 participants showed a significant correlation between ITPC and power. While these single subject-level analyses can show specific instances when the ITPC and power effects co-varied due to subject specific differences or random variability, they do not capture the trends across the entire group. Using linear mixed-effect models that pool all electrodes included in the study, we modeled the correlation between pre-auditory-onset theta ITPC and post-auditory-onset theta power, with *subject* as the random variable. Across all electrodes in the STG, no relationship between power and ITPC effects was observed [*F*(525) = 0.06, *p* = 0.790]. Furthermore, restricting the model to electrodes based on anatomical region showed no significant relationships: anterior STG [*F*(132) = 0.963, *p* = 0.328], middle STG [*F*(215) = 1.095, *p* = 0.297], posterior STG [*F*(174) = −1.757, *p* = 0.071]. These results further reinforce the conclusion that pre and post speech onset effects stem from separate neuronal populations, and likely different mechanisms.

### Comparison of Theta ITPC and Theta Power in Congruent-Audiovisual and Incongruent-Audiovisual Trials

The above analyses test the main proposed hypothesis: that congruent audiovisual ITPC and power effects are largely independent. A comparison of congruent and incongruent audiovisual speech allows a follow-up examination of how theta ITPC and power are modulated when the visual evolution of the word differs from the auditory envelope of the word. Specifically, stimuli were designed so that incongruent visual words nevertheless predicted the same onset timing of the mismatched auditory word, meaning that congruent and incongruent audiovisual stimuli should not differ before sound onset in either ITPC or power measures. Accordingly, a comparison of ITPC and power differences after speech onset enable a better interpretation of the functional significance of the two physiological responses. To this end, we compared the time courses of ITPC for congruent-audiovisual and incongruent-audiovisual speech conditions, similar to the analysis performed between auditory-alone and congruent-audiovisual. The analysis was performed by pooling electrodes within the anterior, middle and posterior STG within each subject, and modeled both the slope and intercept using a linear mixed-effects model.

As shown in Figure 6, for both congruent-audiovisual and incongruent-audiovisual conditions, theta band ITPC increased steadily after the onset of visual mouth movements and peaked quickly after sound onset. As expected, no significant differences in ITPC values were observed in response to visual speech onset. However, audiovisual congruency did introduce a significant difference in ITPC values in the anterior STG (−0.07 to 0.04 secs and 0.26 to 0.5 secs, minimum *p*= 0.015) generally following sound onset (the small differences present before sound onset are likely attributable to temporal smoothing due to spectral filtering). No significant ITPC differences were observed in the middle STG (minimum *p* = 0.999) or posterior STG (minimum *p* = 0.996) at any time point in the ITPC measure.

**Figure 6.**
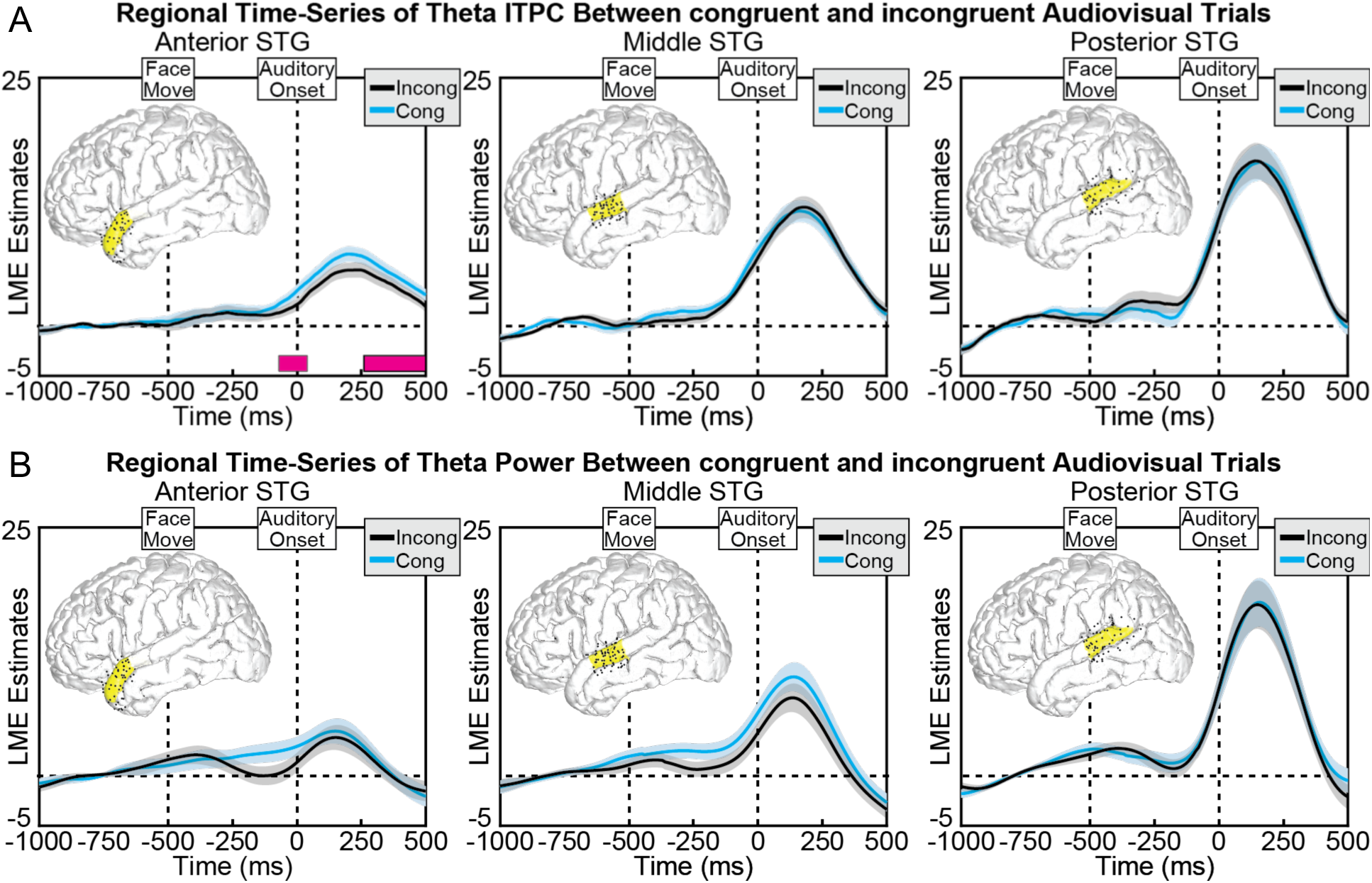
(a) Group linear mixed-effect model (LME) estimates for each time-point of theta theta ITPC in congruentaudiovisual(blue) and incongruent-audiovisual (red) trials, calculated separately at anterior (left), middle (middle), and posterior (right) regions of the STG. Shaded areas reflect 95% confidence intervals. No Significant differences in theta ITPC emerged briefly after the onset of the auditory stimuli, during the time course of the stimuli in the anterior STG. (b) LME estimates for each time-point of theta power in congruent-audiovisual(blue) and incongruent-audiovisual (red) trials, calculated separately at anterior (left), middle (middle), and posterior (right) regions of the STG. Shaded areas reflect 95% confidence intervals. No Significant differences in theta power emerged during the time course of the stimuli.

Following a similar pattern to the ITPC data, theta band power in both congruent-audiovisual and incongruent-audiovisual conditions increased steadily after mouth movement and peaked after auditory onset. Importantly, despite the differences in ITPC values between the two conditions, no significant differences were observed in theta band power in any of the three ROIs, anterior STG (minimum *p* = 0.531), middle STG (minimum *p* = 0.558) and posterior STG (minimum *p* > 0.99).

## Discussion

Visual speech improves speech perception fluency, but the kinds of information that visual speech provides to the auditory system remains unclear. While prior studies have separately demonstrated that visual speech can modulate timing processes both before and after auditory speech onset, it remains unknown whether these processes are enabled by overlapping or independent neural mechanisms. In the current study, we recorded intracranial EEG data from participants as they listened to audiovisual or auditory-alone speech while performing a word detection task. Replicating prior work, we found evidence for visual modulation of speech both before and after speech onset in the STG, but crucially showed a dissociation between the two. These data are consistent with a model in which speech onset information and envelope tracking from the visual stream are encoded by two distinct mechanisms in the STG.

### Correlated and Complementary Modes of Audiovisual Speech Processing

Our findings suggest two distinct mechanisms at work, each associated with specific stages of speech processing. The first mechanism encodes predictive timing information, operating via theta phase reset during preparatory mouth movements and before auditory speech onset in the anterior and middle STG. This phase resetting enables alignment of neural activity to optimize processing in anticipation for the onset of speech. The second mechanism encodes correlated visual speech information during continuous speech in the form of increased theta band power when audiovisual speech is present.

Our findings align with the framework proposed by Campbell (2007), which suggests that visual speech operates in two modes: a “complementary mode” that provides predictive timing cues, such as mouth movements before speech onset, and a “correlated mode” where visual and auditory cues synchronize during ongoing speech. In the “complementary mode”, preparatory mouth movements deliver critical timing information for auditory speech onset; however, once auditory input begins, visual cues become less crucial, as auditory information takes precedence. This interpretation aligns with our observation that visual speech modulates the predictive period before speech onset while exerting a reduced influence afterward.

### Preparatory Mouth Movements Modulate Theta ITPC across the STG

Preparatory mouth movements led to increased theta ITPC across all STG sub-regions, without inducing changes in theta power. This finding replicates evidence that visual cues from mouth movements can reset theta phase in auditory areas (Biau et al., 2021; Mégevand et al., 2020), but with a larger sample of electrodes and patients, as well as using robust group-level statistics. This phase reset appears to contribute to predictive timing of speech onset, potentially optimizing the phase of auditory neuron activity for effective speech onset processing (Arnal et al., 2009; Karas et al., 2019). Indeed, studies suggest that the precise theta phase of auditory neurons can impact visual speech perception outcomes (Thézé et al., 2020), supporting a model in which theta phase reset in response to mouth movements facilitates optimal phase angles for processing auditory speech onset. Moreover, the observed ITPC differences in middle and posterior STG align with auditory speech studies indicating that these regions are functionally specialized for encoding speech onsets (Hamilton et al., 2018).

### Sustained Theta ITPC Differences During Speech Envelope Encoding

We observed heightened ITPC in the congruent-audiovisual condition compared to auditory-alone in the anterior STG, particularly aligned with the auditory speech envelope. Recent evidence from auditory speech studies, such as Oganian et al. (2023), suggests that theta phase resetting occurs primarily as an evoked response to speech onset, rather than as a sustained process throughout speech. Therefore, the ITPC differences observed during the speech envelope may be unique to audiovisual speech processing. However, comparing congruent-audiovisual and auditory-alone conditions alone limits our ability to determine whether these ITPC differences are due to broader attentional processes associated with the presence of a speaker or if they reflect more specific responses to the speech envelope itself.

However, if ITPC differences after speech onset indeed reflect attentional differences, then we would observe much weaker ITPC differences during the speech envelope between congruent-audiovisual and incongruent-audiovisual. That is not what we found. Differences between congruent-audiovisual and incongruent-audiovisual speech were significant after speech onset, suggesting that ITPC in the anterior STG may capture aspects of envelope tracking. Indeed, previous research has indicated that the anterior STG is functionally specialized for sustained speech encoding, as seen in its tracking of the speech envelope through high gamma power (HGp; a correlate of population spiking rates) (Hamilton et al., 2018).

Recent findings have suggested that HGp is modulated by theta oscillatory phases during audiovisual speech perception (Mégevand et al., 2020). Considering the sustained theta ITPC differences observed in the anterior STG, our results may offer a mechanistic basis the sustained speech encoding observed in HGp by Hamilton et al., 2018. This theta-to-HGp coupling could represent an adaptive process in the anterior STG, optimizing speech envelope tracking and potentially facilitating the integration of multisensory cues for more effective audiovisual speech processing.

### Theta Power and Attention Modulation in Posterior STG

In addition to prominent ITPC differences, we observed theta power differences in the posterior STG (pSTG) after speech onset between the congruent-audiovisual and auditory-alone conditions, but not between the congruent-audiovisual and incongruent-audiovisual conditions. These findings suggest that the observed theta power differences in pSTG are not reflecting speech envelope related information in the visual stream, but rather processes that are different only between auditory-alone and congruent-audiovisual. Thus it is possible that theta power differences in pSTG are reflecting attentional modulation tied to the presence of multisensory input rather than processes unique to speech envelope encoding.

These findings are consistent with recent work suggesting that theta oscillations play a role in modulating attention during audiovisual speech. Specifically, Byczynski and Park (2024) demonstrated that faster theta rhythms coordinate attention by suppressing irrelevant input to optimize focus on relevant stimuli. Wang et al. (2016) further showed that theta oscillations engage cognitive control networks during cross-modal attention tasks, supporting a flexible and focused attentional state when congruent audiovisual cues are present. Together, our findings and these studies indicate that theta power in the posterior STG likely plays a broader attentional role, enhancing attention allocation in response to congruent visual and auditory signals.

While some studies have shown that congruent mouth movements aid auditory speech encoding significantly better than incongruent speech in broadband ERP signals (e.g. Crosse et al., 2015), which is largely represented by low-frequency signals including theta activity, the absence of theta power differences in our study does not contradict these findings. As broadband ERP signals reflect a composite of multiple frequency bands, it is possible that other frequencies, including delta activity which has been suggested to reflect comprehension level encoding of the speech envelope (Molinaro & Lizarazu, 2018), contributes to the envelope tracking differences seen in studies like Crosse (2015).

### Limitations

While these findings offer valuable insights, the generalizability of our study is limited by the scope of stimuli used. Our analyses focused on whole-word stimuli grouped by four specific phonemes (/b/, /d/, /g/, /h/), raising the possibility that the observed effects may not apply to a broader range of consonants. Additionally, our use of word level stimuli, while optimal to capture pre-auditory-onset and post-auditory-onset effects for the study, does not reflect sentence level dynamics observed in more naturalistic stimuli seen in other studies (e.g. Oganian et al., 2023, Mégevand et al., 2020).

### Conclusion

In summary, the current study found pre-speech onset theta ITPC differences across the STG, as well as post-speech onset theta power differences, operating via two separate mechanisms. We replicate and expand on pre-speech-onset ITPC effects (Mégevand et al., 2020) as well the post speech onset theta band power differences in the posterior superior temporal gyrus (Karthik et al., 2021), providing a unified analysis of these separate processes. Together, these findings point to a dual mechanism in audiovisual speech processing: one that encodes predictive timing via phase reset and another that encodes speech correlation via sustained theta activity. Importantly, our data show that phase and power encoding are independent and are processed by largely non-overlapping neural populations within the STG.

## Notes

### Competing Interest Statement

The authors have declared no competing interest.

