## Supplemental Information for "Visual speech enhances auditory onset timing and envelope tracking through distinct mechanisms"

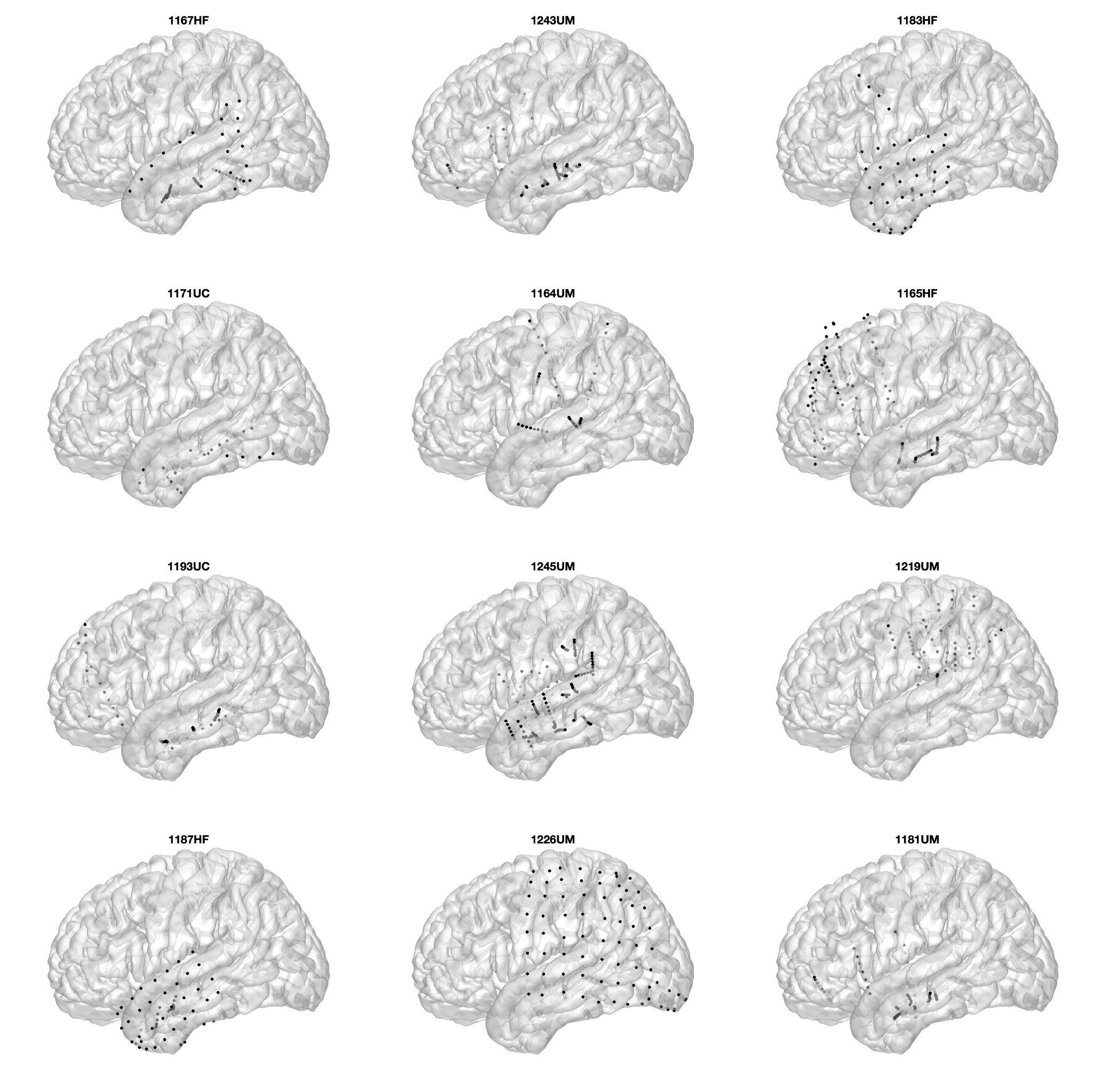

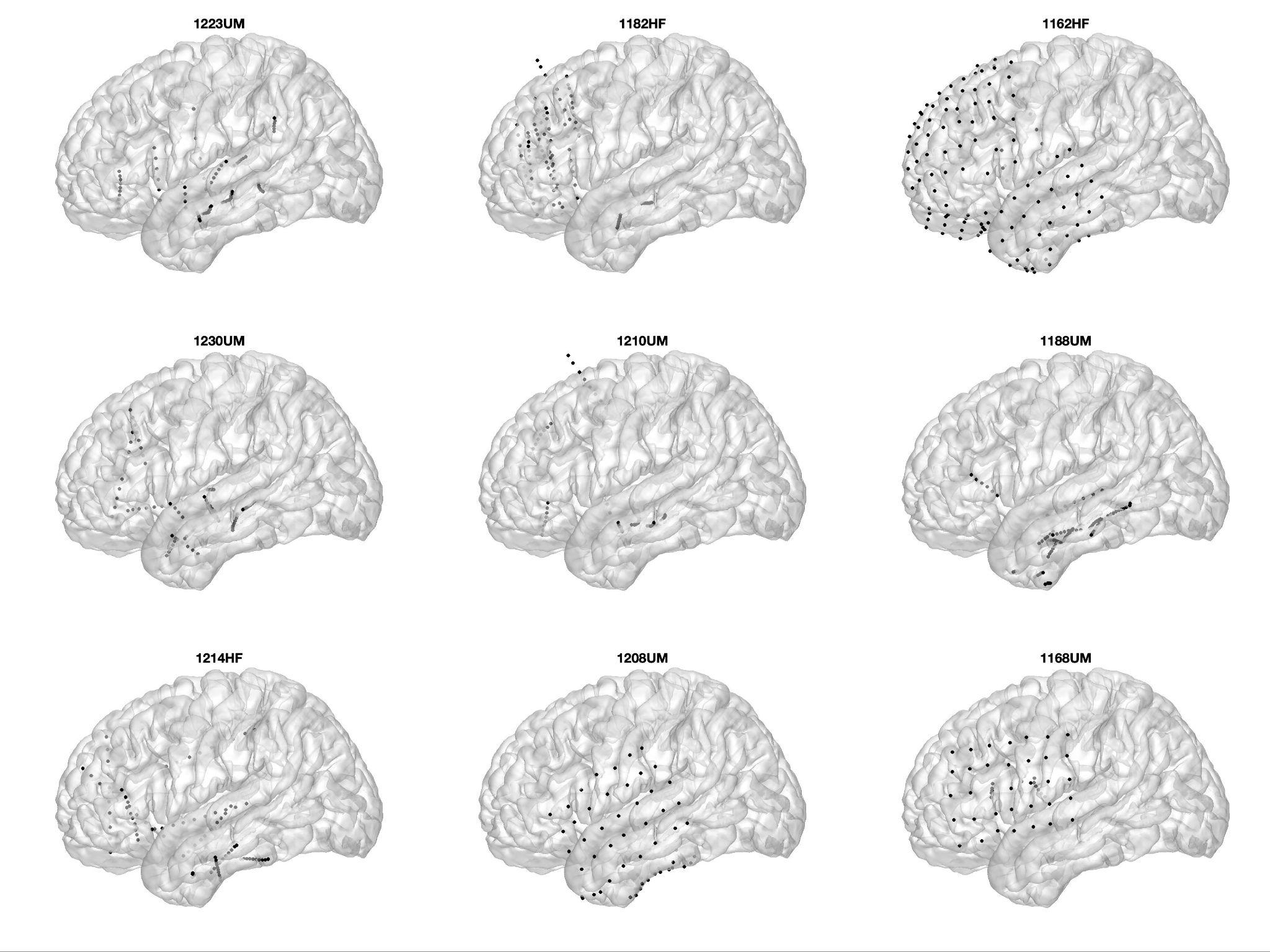

***SI Figure 1***, Electrode coverage of participants in the study. 21 subjects, N= 1664 total. Only electrodes that met anatomical selection criteria (located within the superiortemporal gyrus, or STG) and functional selection criteria were considered.

**SI Table 1: Word stimuli used in the study**

| bag | dad | fad | gag |
| --- | --- | --- | --- |
| bank | dance | fast | gang |
| base | dash | fat | gas |
| beard | dial | file | gear |
| beer | digs | fill | gig |
| bias | dill | fine | gill |
| bible | dine | fish | guild |
| bid | dish | fist | guile |
| bill | disk | fit | guise |
| bye | dive | five | guy |
